# A macroevolutionary role for chromosomal fusion and fission in *Erebia* butterflies

**DOI:** 10.1101/2023.01.16.524200

**Authors:** Hannah Augustijnen, Livio Bätscher, Martin Cesanek, Tinatin Chkhartishvili, Vlad Dincă, Giorgi Iankoshvili, Kota Ogawa, Roger Vila, Seraina Klopfstein, Jurriaan M. de Vos, Kay Lucek

## Abstract

The impact of large-scale chromosomal rearrangements, such as fusions and fissions, on speciation is a long-standing conundrum. We assessed whether bursts of change in chromosome numbers resulting from chromosomal fusion and fission are related to increased speciation rates in *Erebia*, one of the most species-rich and karyotypically variable butterfly groups. We established a genome-based phylogeny and employed state-dependent birth-death models to infer trajectories of karyotype evolution across this genus. We demonstrated that rates of anagenetic chromosomal changes (*i.e*. along phylogenetic branches) exceed cladogenetic changes (*i. e*. at speciation events), but when cladogenetic changes occur, they are mostly associated with chromosomal fissions rather than fusions. Moreover, we found that the relative importance of fusion and fission differs among *Erebia* clades of different ages, where especially in younger, more karyotypically diverse clades, speciation is more frequently associated with chromosomal changes. Overall, our results imply that chromosomal fusions and fissions have contrasting macroevolutionary roles and that large-scale chromosomal rearrangements are associated with bursts of species diversification.

## Introduction

The evolution of barriers to gene flow is a critical requirement for the progress of speciation [^1^]. Although several barriers may contribute to the process, their relative importance often remains unknown, especially at a macroevolutionary scale [^2^]. Chromosomal speciation theory suggests that large-scale chromosomal rearrangements (CRs), such as fusions and fissions, are able to promote speciation. They may allow for the build-up of genetic incompatibilities between lineages either by causing hybrid dysfunction [^3,4^] or by suppressing recombination in rearranged sections of the genome [^5,6,7^]. The relevance of chromosomal speciation has been criticized due to the expected “underdominance” of CRs; whereby strong hybrid fitness disadvantages ensure that fixation of novel karyotypes is difficult, rendering barrier formation unlikely [^5,6^]. Paradoxically, if the effects of CRs on hybrids were minor, fixation would be possible, but the resulting barriers would remain shallow [^5,6^]. Importantly, these theories were developed for monocentric chromosomes, whereas the chromosomes of several major organismal groups, such as butterflies and sedges [^8^], are holocentric, *i.e*. they have centromere-like structures spread across their chromosomes rather than concentrated in a single centromere [^9^].

Holocentric chromosomes may be more likely to overcome the aforementioned underdominance paradox, as rearranged chromosomes can retain kinetochore functionality and segregate correctly in hybrids [^9,10,11^]. Crosses between closely-related holocentric species with different karyotypes may remain viable [^10^] and do not necessarily result in reproductive isolation [^11,12^]. In addition, some holocentric clades have evolved mechanisms to facilitate proper chromosome segregation even when chromosomes are rearranged [^9,13,14^], which has been suggested to promote chromosomal speciation [^9^]. Empirical evidence for a link between speciation and CRs, especially for chromosomal fusions and fissions, is sparse for both mono- and holocentric clades [^8,15^]. But the fact that many holocentric groups within plants and invertebrates are very species-rich suggests that CRs could have driven diversification in some of them [^9,16^].

Lepidoptera is one of the largest taxonomic groups with holocentric chromosomes, comprising more than 160’000 species of butterflies and moths [^17^]. While some clades within Lepidoptera are extremely diverse in chromosome numbers, sometimes differing by a count of more than 200 even within a single genus [^18,19^], most others have conserved chromosome numbers, often close to the inferred ancestral karyotype (n = 31) [^20,15^]. Comparative phylogenetic analyses indicate a positive association between the rate of speciation and karyotype evolution for several of the most karyotypically diverse butterfly genera [^15^].

*Erebia* is one of the most speciose of all Palearctic butterfly genera, consisting of around 90-100 species that mainly inhabit cold mountainous regions, with the majority of diversity found in Europe, where closely related species often form narrow contact zones [^21^]. Notably, *Erebia* is also one of the genera with the highest karyotype diversity among butterflies [^22^], although this diversity differs between clades within the genus. Most of the variation can be found in the comparatively young *tyndarus* clade (n = 8-51) (Fig. 1B, Table S1), where phylogenetic relationships have remained unclear [^23,24^]. Here, we leveraged karyotype diversity across *Erebia* to test for its role in species diversification. Specifically, we first quantified the overall impact of chromosomal fusion and fission on diversification in *Erebia* using phylogenomic inference and Bayesian state-dependent birth-death models. We then assessed the association between diversification and chromosomal fusion and fission across *Erebia* clades of different ages and karyotype diversity. We hypothesized that chromosomal speciation has played a significant role in the diversification of *Erebia* and that younger, more karyologically diverse clades show an increased signal of chromosomal changes and higher associated speciation rates.

**Figure 1:**
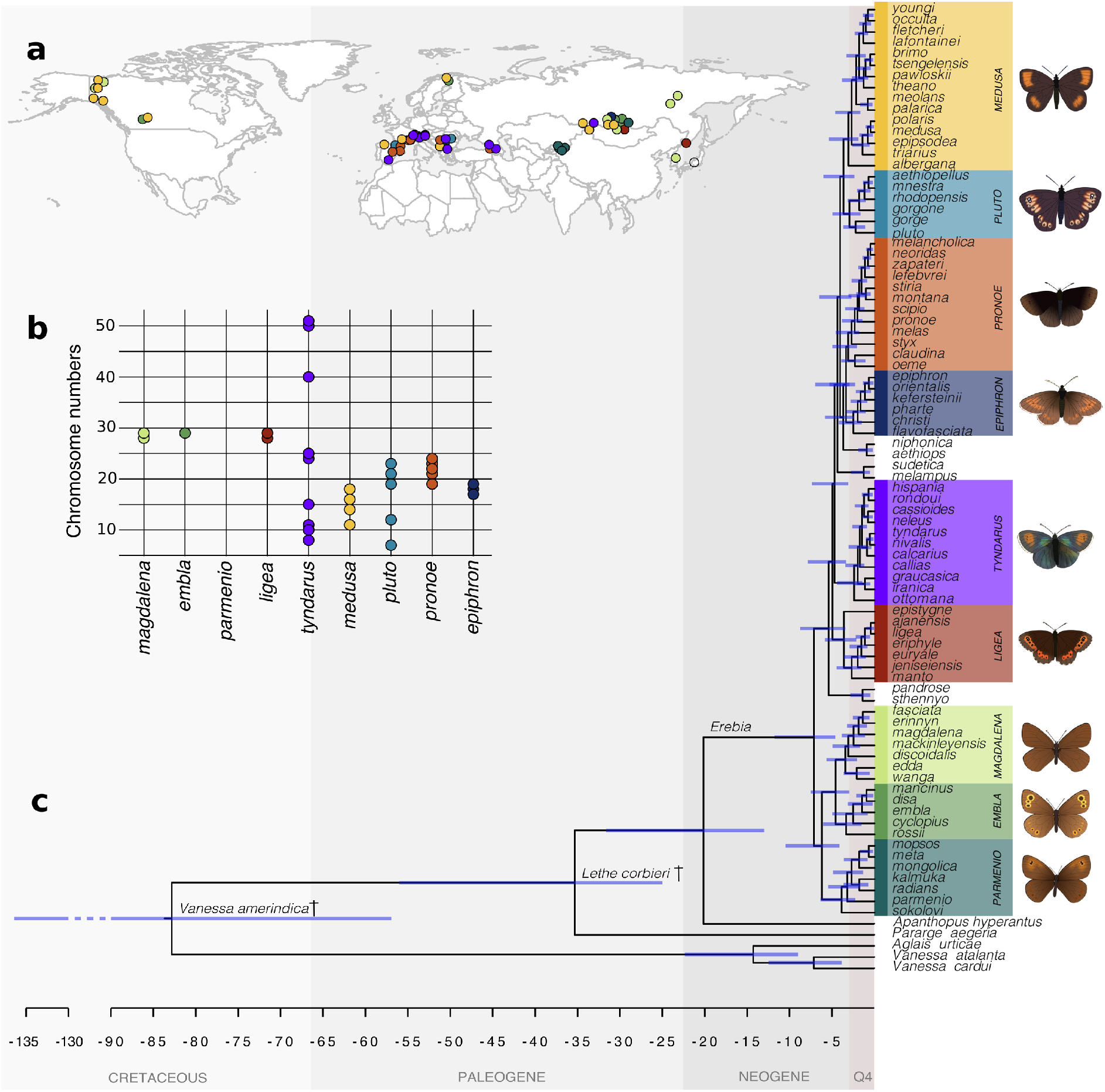
Sample distribution and relationships within the Palearctic genus Erebia. A. Map of the Northern Hemisphere indicating sampling locations of Erebia specimens used, coloured by clade. B. Known chromosome numbers of Erebia species, grouped by clade (from ^21^). C. Time-calibrated phylogeny of Erebia calculated in MrBayes. The fossil Vanessa amerindica was used as a calibration point at the root of the tree, while Lethe corbieri was placed at the base of Satyrini. Clades are based on [^21^]with the exception of medusa and pluto. For each clade a representative phenotype is shown.

## Results

### *The diversification of* Erebia

Although approximately 90-100 *Erebia* species are recognized, the phylogenetic resolution of former studies on *Erebia* was limited, especially for evolutionary younger taxa, as they included either few genes [^21^] or few species [^24^]. Using whole-genome resequencing data for 82 *Erebia* species, (Fig. 1A-B, Table S1-2), we constructed a nearly fully resolved phylogeny based on coalescence of 2’920 individual maximum-likelihood gene trees (Fig. S1). Branch support was very high overall (>0.9 ASTRAL consensus for 96.5% of all nodes), and we further validated the relationships among taxa following [^25^] (Fig. S2-S6). We used the resulting topology to constrain a dating analysis in MrBayes based on two calibration points, using a subset of genes selected for minimal missing data, especially among outgroups (Figure 1C). We confirmed the monophyly of previously defined [^21^] clades *tyndarus* (2.41 Myr, 95%HPD: 1.18-3.43 Myr), *epiphron* (2.47 Myr, 95%HPD: 1.49-4.21 Myr) and *pronoe* (3.10 Myr, 95%HPD: 2.07-4.98 Myr), as well as the classic taxonomic clades [^26^] *ligea* (3.60 Myr, 95%HPD: 2.14-5.81 Myr), *medusa* (3.08 Myr, 95%HPD: 1.87-5.38 Myr) and *pluto* (2.94 Myr, 95%HPD: 1.64-4.93 Myr). We estimated the age of *Erebia* to be 20.16 Myr (95% highest posterior density (HPD) interval: 12.98-31.63 Myr), with the first major split between the mostly non-European *embla*, *magdalena* and *parmenio* clades and all other *Erebia* at 7.14 Myr ago (95%HPD: 4.62-11.77 Myr).

### Cladogenesis and chromosomes

We fitted a ChromoSSE model [^27^] to decompose rates of chromosomal fusion and fission into their anagenetic (chromosomal change along a branch) and cladogenetic (chromosomal change at a speciation event) components. We found that the majority of chromosomal changes occurred by anagenetic fusion (0.631 Events/Species/Myr, 95%HPD = 0.151-1.119) and to a lesser degree fission (0.213 Events/Species/Myr, 95%HPD: 0.003-0.547, Fig. 2). However, most inferred speciation (cladogenetic) rates in *Erebia* coincide with chromosomal change, either with cladogenetic chromosomal fusion (0.197 Events/Species/Myr, 95%HPD: 0.079-0.354) or fission (0.328 Events/Species/Myr, 95%HPD: 0.064-0.563, Fig. 2). Cladogenetic speciation rates under a “no chromosomal change” scenario were lower (0.128 Events/Species/Myr, 95%HPD: 0.087-0.326). The relative extinction rate across *Erebia* was 0.273 Events/Species/Myr (95%HPD = 0.001-0.316) and total speciation (cladogenetic speciation rates for fusion, fission, and without chromosomal change combined) summed to 0.653 Events/Species/Myr (95%HPD: 0.474-0.832).

**Figure 2:**
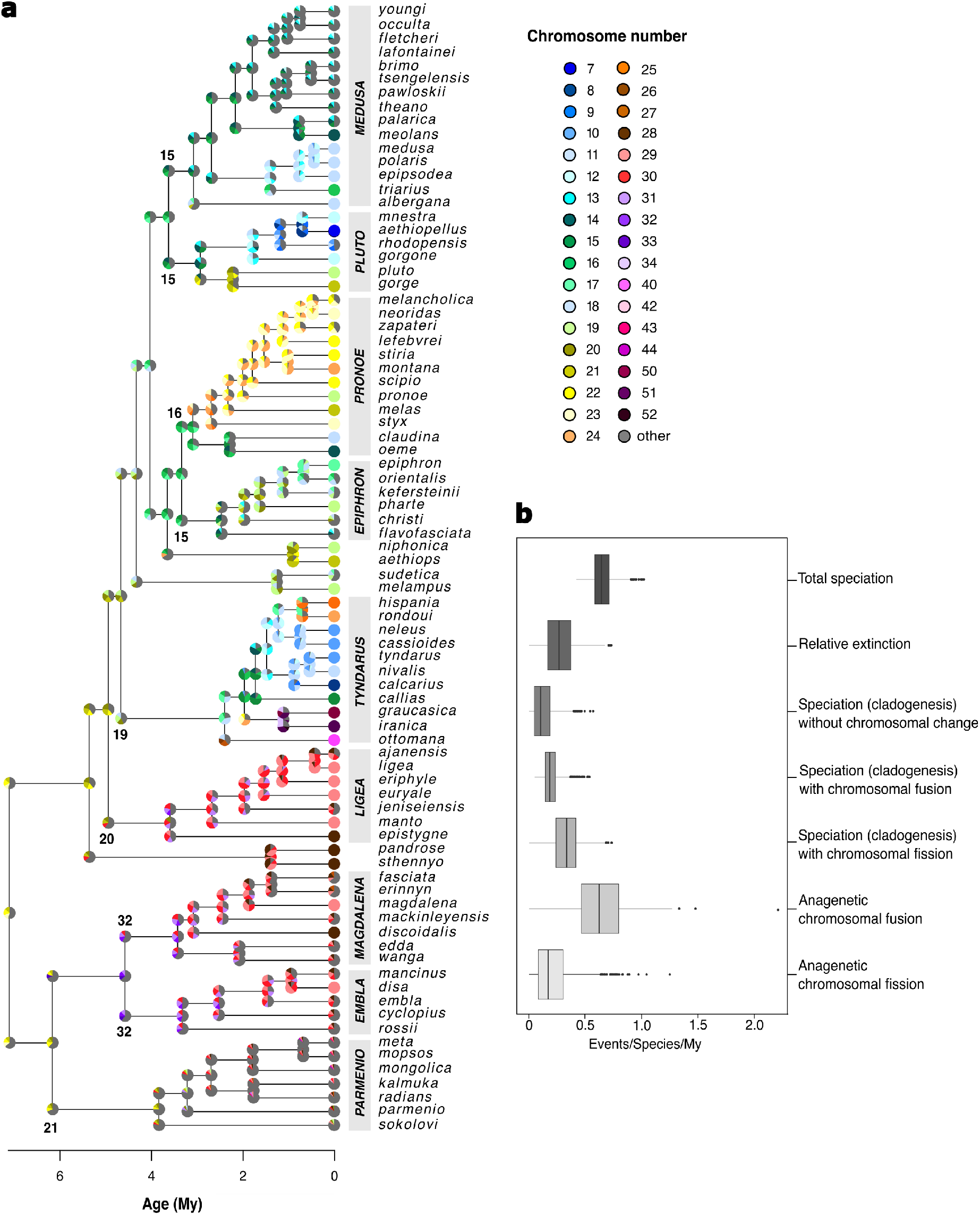
Summary of the chromosome evolution model for Erebia, implemented in ChromoSSE [^27^] A) Estimated ancestral chromosome numbers for Erebia, inferred using the phylogenomic topology of Fig. S1. Chromosome numbers are indicated proportionally by the colour of the pie charts at the branch nodes. The pie charts at the “shoulders” of each node represent the inferred chromosomal state immediately after a speciation event. The dominant inferred chromosome number of the ancestor of each clade is depicted atop or below the node that starts that clade. Clades are named as in Fig. 1. B) Summary of the posterior distributions of inferred ChromoSSE parameters for the overall Erebia analysis, also listed in Table S3.

### Background speciation

State-dependent speciation and extinction (SSE) models, including ChromoSSE, may be prone to Type I errors because shifts in diversification rates across a phylogeny could be assigned to a studied trait even when such shifts are instead caused by an undetected and unrelated trait [^28^]. We therefore estimated diversification rates independently of chromosome states for *Erebia*, to assess whether such underlying speciation rate heterogeneity may influence our inference of ancestral chromosomal changes and rate shifts. We employed MiSSE [^29^] an extension of the “hidden-states” state-dependent model HiSSE [^30^] that allows to model background diversification resulting from hidden causes of speciation. Our inference indicated that while there seem to be some shifts in diversification rates along terminal branches, there is an apparent lack of diversification shifts at deeper parts of the tree (Fig. 3). Our ChromoSSE inference, especially of cladogenetic parameters at the nodes, is therefore unlikely to have been affected by such background speciation. If hidden speciation had any influence at all, it would be a minor increase of estimates of recent anagenetic chromosomal changes in those terminal branches that do show small speciation rate shifts.

**Figure 3:**
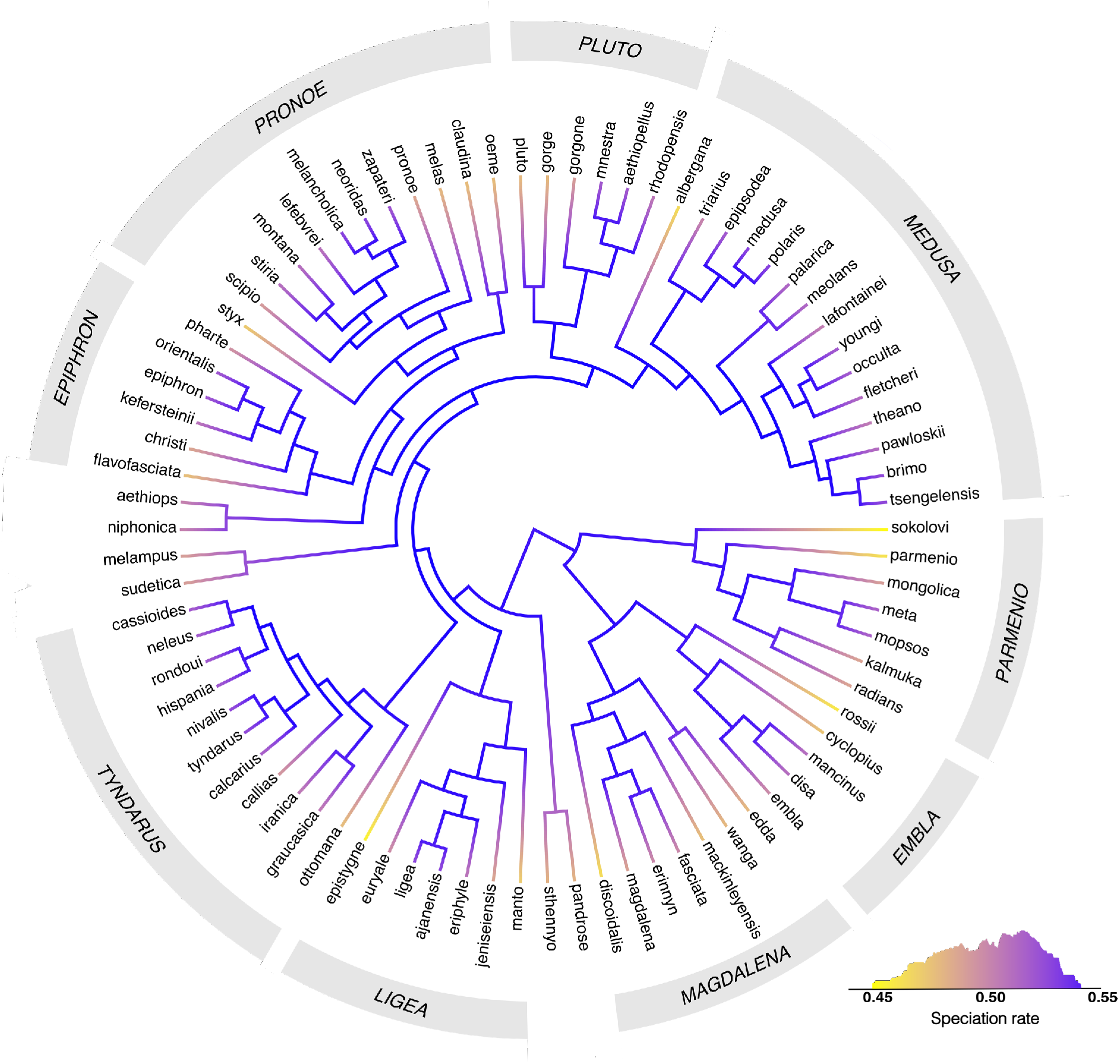
“Hidden States” trait-independent inference of speciation rates across the Erebia phylogeny, computed using MiSSE [^29^]. Colors indicate inferred speciation rates based on model averaging of the 10 best models as selected by the greedy MiSSE algorithm from 28 possible combinations of hidden state rates (see Table S6). Hidden states are estimations of factors that could influence speciation in the “background” and thus cause diversification shifts which may negatively impact an SSE inference. Speciation rates generally show a decrease only in the terminal branches, whereas internal nodes show no major shifts.

Focusing only on the tips of the phylogeny, we showed that background speciation rates at the tips ranged from 0.446 to 0.526 (mean 0.496 ±0.021; Fig. 3) and differed between clades (ANOVA: *F*_8,67_ = 2.50, *p* = 0.020; Fig. S7), where *tyndarus* (0.510 ±0.015) and *medusa* (0.510 ±0.016) showed the highest rates. Current background extinction rates also differed among clades (*F*_8,67_ = 2.38, *p* = 0.026, Table S2) and were lower than speciation rates, ranging between 0.003 and 0.012 (mean 0.006 ±0.002), where the lowest extinction rates occurred in the clades that showed the highest speciation rates (Fig. S7).

### Clade-specific chromosomal changes

To test whether ana- or cladogenic chromosomal change rates would be higher in clades that show a higher karyotype diversity (Fig. 1B), we ran ChromoSSE for each clade for which we had sufficient chromosome count data (Fig. 4A). We found that the rates of both ana- and cladogenetic chromosomal change differed across clades under ChromoSSE: Anagenetic fusions differed significantly among clades (ANOVA: *F*_5,14994_ = 847.9, *p* < 0.001; Fig. 4B), with posterior distribution means ranging from comparatively low: 0.084 (*ligea*), to moderate: 0.280 Events/Species/Myr (*pronoe*). This was also true for anagenetic fissions (*F*_5,14994_ = 409.3, *p* < 0.001; Fig. 4C), with mean rates between 0.065 (*ligea*) and 0.198 Events/Species/Myr (*pronoe*). Total anagenetic chromosomal change, defined as the sum of rates for anagenetic fusions and fissions, ranged from 0.148 (*ligea*) to 0.478 Events/Species/Myr (*pronoe*), which for all clades is less than the 0.844 (95%HPD = 0.420-1.295) found for the overall ChromoSSE analysis of *Erebia* (Table S3).

**Figure 4:**
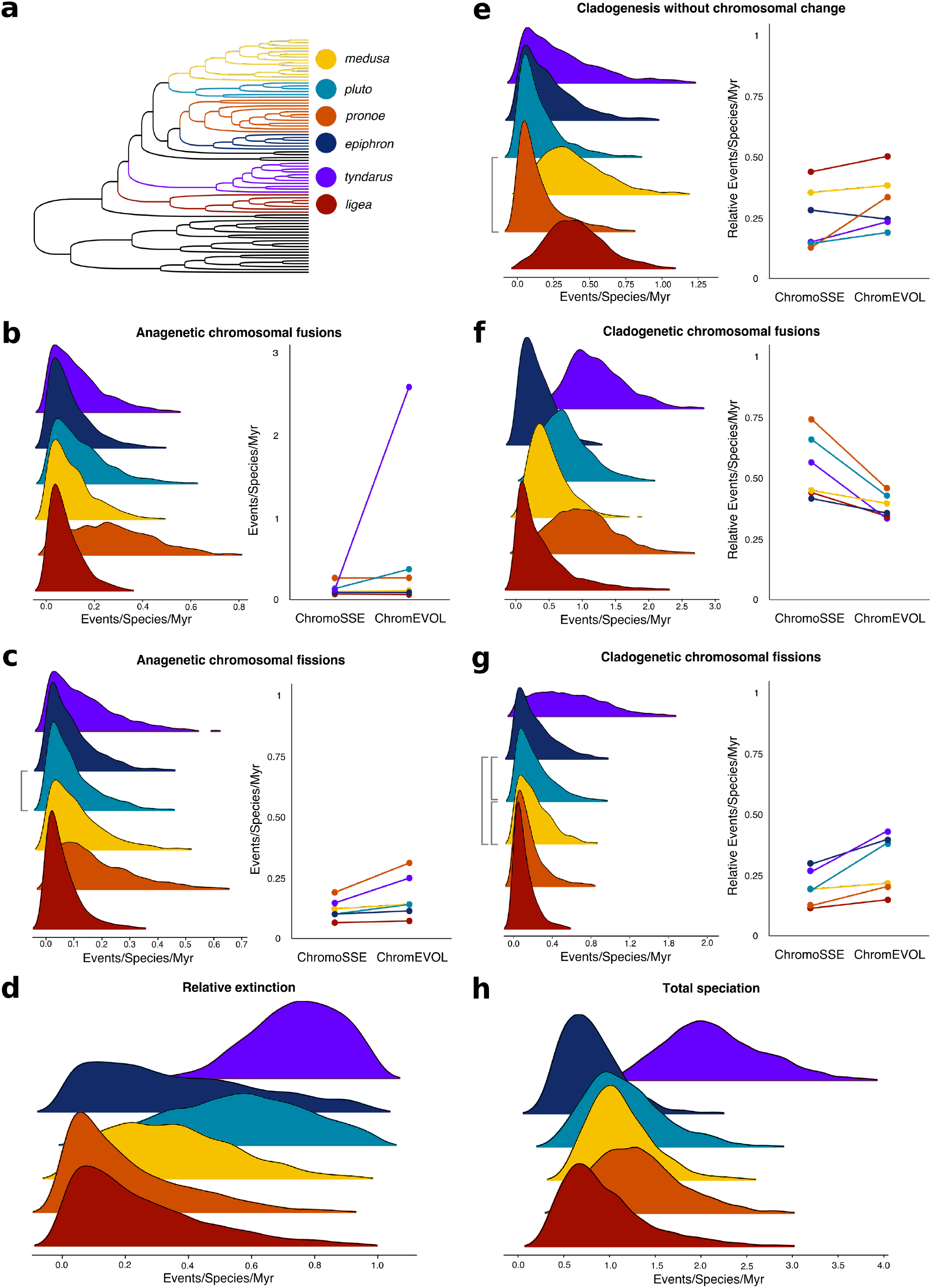
Summary of ana- and cladogenetic ancestral chromosome estimations for six Erebia clades A) Overview of the phylogeny of Erebia with relevant clades highlighted. B-H) Distributions of the posterior probability estimates for: B) anagenetic fusions C) anagenetic fissions D) relative extinction rates E) cladogenetic changes unrelated to chromosomal change F) cladogenetic chromosomal fusions G) cladogenetic chromosomal fissions H) total speciation rates. For B-C and E-G comparisons between standardised parameter estimations of clade specific state-dependent birth-death models (ChromoSSE) and birth-death independent models (ChromEVOL) are shown. Each dot indicates the mean of the posterior probability space. All parameter rates are expressed in events per species per million years. Grey connecting bars denote clades whose posterior probability estimates do not differ in a post hoc comparison (all others p<0.001).

Rates for speciation (cladogenesis) with chromosomal fusion showed a higher degree of differentiation among clades than for other ana-or cladogenetic parameters (*F*5,14994 = 1908.0, *p* < 0.001; Fig. 4F), ranging from 0.328 (*epiphron*) to 1.263 Events/Species/Myr (*tyndarus*). Rates for speciation (cladogenesis) with chromosomal fission were likewise variable between clades (*F*5,14994 = 1282.0, *p* < 0.001; Fig. 4G), ranging between 0.115 (*ligea*) and 0.594 Events/Species/Myr (*tyndarus*). Total speciation (cladogenesis) with chromosomal change (the sum of cladogenetic fusions and fissions) ranged from 0.537 (*ligea*) to 1.856 Events/Species/Myr (*tyndarus*). These rates were generally higher than for overall *Erebia* (0.528 Events/Species/Myr), but the posterior distributions for most clades overlap with the 95%HPD (0.304-0.783) of the overall parameter (Table S3). The exception was the karyotypically diverse *tyndarus* clade (95%HPD = 0.906-2.898). While the parameter values were very similar between some clades, their proportional contributions to rates of speciation differ (Fig. 4). For example, *medusa, pluto* and *epiphron* show similar rates of cladogenetic chromosomal fissions but they account for 19.8%, 19.8% and 30.3 % of the total cladogenetic events, respectively.

Speciation rates (cladogenesis) without chromosomal change differed among clades (*F*_5,14994_ = 655.3, *p* < 0.001; Fig 4E) ranging from 0.167 (*pronoe*) to 0.421 (*ligea*) Events/Species/Myr, but never accounted for a majority of the total speciation in a clade, though in *ligea* it nearly equalled the degree of cladogenetic fusions (Fig. 4E, F; Table S3).

Relative extinction rates were highest for *tyndarus* at 0.745 and varied between clades (*F*_5,14994_ = 2432.0, *p* < 0.001; Fig. 4D), with the lowest rates for *pronoe* (0.211) and *ligea* (0.251). Total speciation related directly to rates of cladogenetic events, and likewise varied between clades (*F*_5,14994_ = 2737.0, *p* < 0.001; Fig. 4H), with *tyndarus* having the highest degree of total speciation (2.191, 95%HPD = 1.283-3.363 Events/Species/Myr). The correlation between relative extinction and total speciation was 0.38 (*t_1,14998_* = 50.7, *p* < 0.001).

### A tale of two models

In order to assess to which degree the combined inference of speciation and extinction rates in ChromoSSE might affect our estimations of cladogenetic and anagenetic parameters, we validated the results of ChromoSSE by carrying out similar analyses in ChromEVOL [^31^] following [^27^], which allows to model cladogenetic changes without extinction. We expected potential differences between the models in clades with higher extinction and speciation rates. Although the results of ChromoSSE and ChromEvol were largely similar for nearly all clades (Table S4, Figs. 4, S8) the marginal likelihoods of ChromEvol models were generally higher. The difference in likelihood was more pronounced for the overall tree than for individual clades, where likelihood differences were not as high (difference in Bayes Factors (BFs) ranging from to 10.762 (*pluto*) to 39.433 (*medusa*); Table S4). In contrast to our predictions, we found no significant difference between model fits (BF = 1.078) for *tyndarus*, the clade with the highest inferred extinction rates (Fig. 4D). An explanation for the difference between ChromoSSE and ChromEvol could be that simpler models with fewer parameters (ChromEvol) result in higher marginal likelihoods than more complex models which may suffer from overparameterization, particularly in relatively small datasets. In clades with low to medium inferred relative extinction rates (*e.g. medusa*) the more complex ChromoSSE model was not preferred over the relatively simpler ChromEvol. In *tyndarus*, by contrast, it appears that modelling extinction in tandem with chromosomal evolution as ChromoSSE was designed to do, was sufficient to overcome the potential issue of overparameterization.

Since ChromEvol estimates cladogenetic parameters as proportions of the total amount of cladogenetic change, cladogenetic parameters from the ChromoSSE analysis were divided by the average total speciation (Fig. 4G) allowing us to compare relative Events/Species/Myr. For the entire tree, rates of cladogenesis without chromosomal change, cladogenetic fusions and cladogenetic fissions overlapped in their posterior distributions, but posterior means differed between the models (*t*-test, all *p* < 0.001, Fig. S8), as would be expected given the relatively high difference between Bayes Factors. However, rates of speciation (cladogenesis) with chromosomal fission remained higher than those for chromosomal fusion under both models. Rates for anagenetic fissions showed a higher overlap in posterior distribution, and despite a significant difference in means (*t*_1,1590_ = 2.1, *p* = 0.040), they showed a similar proportion of anagenetic changes under both models (Fig. S8). Rates of anagenetic fusions differed to a higher degree between models (*t*_1,1590_ = 161.2, *p* < 0.001; Fig. S8). This difference was most likely driven by *tyndarus* clade (see Fig. 4B) and may be related to high relative extinction rates.

## Discussion

Compared to other rearrangements, such as inversions, much more remains unknown about the role of chromosomal fusion and fission in speciation [^8^]. Theory predicts that fusions and fissions could have major impacts on the speciation process [^3,6^], particularly in species whose chromosomes are holocentric, and therefore may not initially be as strongly impacted by the underdominance paradox [^8,10,13^]. Here, we found phylogenomic evidence for an association of chromosomal fusion and fission with higher speciation rates at macroevolutionary scale, providing indirect evidence for their involvement during speciation. From a phylogenetic perspective, chromosome-associated speciation should mainly concern cladogenesis, *i.e*. the splitting of an ancestral species into two new lineages. However, recent theory predicts that anagenetic chromosomal changes could also contribute to speciation by gradually building up differentiation in karyotypes over time, potentially leading to increased reproductive isolation [^8^]. Overall, most changes in chromosome number in *Erebia* appear to have occurred anagenetically (Fig. 2). Chromosomal fusion and fission could therefore act as barriers to gene flow, similarly to other rearrangements [^32^], including inversions [*e.g*.^33^]. These could suppress recombination in rearranged sections of the genome and so prevent the breakup of linkage disequilibrium between locally co-adapted genes [^6^], promoting the gradual accumulation of differences through time [^34,35^]. Speciation may then be completed through additional processes [^36^], such as reinforcement upon secondary contact, as has been suggested for other butterflies [^37^]. However, chromosome-associated cladogenetic events were also prevalent in *Erebia* and were associated with higher rates of speciation than speciation without chromosomal change (Fig. 2B). Such cladogenetic events coincide with splits between lineages, and in theory represent cases where chromosomal rearrangements may have a more causative involvement in speciation [^27^]. This may especially be the case when chromosomal rearrangements act as intrinsic barriers to gene flow, *e.g*. by resulting in hybrid dysfunction [^3,4^], by physically together bringing sites under selection [^32^], or if sex chromosomes are involved [^38^]. Chromosomal fusions and fissions may therefore accompany rapid diversification of lineages into species radiations [*e.g*. ^39^], for instance here in *Erebia* [^21^].

Our analyses suggest that the evolutionary impacts of chromosomal fusions and fissions differ, as fusions were more likely to be anagenetic across the entire *Erebia* phylogeny. Conversely, fissions were more likely to be associated with cladogenetic events, leading to higher cladogenetic speciation rates when fissions are involved (Fig. 2). Reductions in chromosome number through fusion events were more common than fissions in *Erebia*, as well as other groups of butterflies [^15^] and holocentric organisms [*e.g*. ^40^]. Chromosomal fissions, though rarer, may be associated with higher speciation rates because they are predicted to more often result in deleterious meiotic multivalents [^15,41^], although this effect may be partially mitigated in Lepidoptera [^13,42^]. Shorter chromosomes have also been shown to have an increased likelihood to be involved in fusions [^43^], suggesting that longer chromosomes may be more evolutionary stable. The apparent higher stability of fused chromosomes and the instability of fissions could explain why clades with higher chromosome numbers show higher cladogenetic speciation rates in *Erebia* (Fig. 2, 4) and other butterflies [^15^]. The higher overall degree of anagenetic fusions observed in this study might reflect their ability to gradually reduce recombination rates, as has been found in mice [^44^], and other butterflies [^45^]. Genome-wide crossing-over rates correspondingly seem to be considerably higher when chromosomes are short, or evolve to become shorter [^46^, but see ^47^].

While there are some examples of studies focusing on the macroevolutionary impact of chromosomal fusions and fissions, as well as other rearrangements [^48,49,50^], understanding patterns of chromosome-associated speciation may require additional analysis at a finer taxonomic scale [^40^]. Here, we assessed the relative contributions of chromosomal fusion and fission between clades of differing ages and karyotype diversity, in order to assess the impact of these rearrangements on speciation. We identified a continuum of chromosome-associated speciation rates, ranging from the young, karyotypically very diverse *tyndarus* clade that showed the highest rates of speciation related to chromosomal change (Fig. 4) to karyotypically more conserved clades, such as *ligea*, where the proportion of speciation unrelated to chromosomal change was much higher (Fig. 4E). Other clades fell along this continuum, where rates of speciation with chromosomal fusion were generally high and anagenesis appears to be less important at the subclade level than for the overall tree (Fig. 4). The high karyotype diversity and associated high speciation rates of the *tyndarus* clade may in part be explained by their ecology. Species of the *tyndarus* clade occur almost exclusively in Alpine areas [^22^], whereas other *Erebia* clades are ecologically more diverse [^23,51^]. During glacial cycles, repeated range expansions and contractions across relatively small geographic areas, have caused population subdivisions of many Erebia species [*e.g*. ^52,53^] including the *tyndarus* clade [^54^]. For the latter, drift and other stochastic processes could have promoted the fixation of novel chromosomal rearrangements [^55^].

While state-dependent speciation and extinction models, *i.e*. SSE type models, allow for unprecedented phylogenetic insights into macroevolutionary aspects of speciation [^29,56,57^], their reliability has been partially questioned [^58^]. They may, for example, suffer from excess false positive rates because shifts in diversification rates across the phylogeny may be assigned to a studied trait in an SSE model even when these shifts are caused by an undetected and unrelated trait [^28,58^]. This could lead to an overestimation of inferred parameters. However, simulations have indicated that ChromoSSE is more likely to underestimate, rather than overestimate, cladogenetic changes, implying that our inferences are instead rather conservative [^27^]. Furthermore, as the model cannot detect “cryptic” chromosomal rearrangements, *e.g*. fusion and fission events that counter balance each other and therefore do not lead to changes in chromosome number, our data may even represent an underestimation of the effects of chromosomal rearrangements on speciation. For example, Mackintosh et al. 2022^59^ found nine fusion and fission events that may have contributed to speciation between two *Brenthis* butterflies that otherwise differ little in their chromosome numbers. We also highlight that a solid taxonomic framework, as presented here, is highly preferable for the correct interpretation of SSE-based inferences. However, given the taxonomic complexity of *Erebia* [^21^], it is likely that some cryptic species may exist, again indicating that our estimates of diversification may be rather conservative.

We further confirmed the validity of our inferences by fitting ChromEvol [^31^] models that do not estimate extinction and may thus not suffer the same potential pitfalls as SSE type models. We found very similar results as for the SSE-based analyses (Fig 2B, 4), with the exception of inferred rates of anagenetic fusions in the *tyndarus* clade, which were estimated to be much higher under ChromEvol. This difference may be due to the influence of the high relative extinction rates in this clade (Fig. 4H), as ChromEvol does not consider any unobserved speciation that may have resulted in extinction, whereas ChromoSSE does [^27^]. Also, by estimating “hidden” background speciation and extinction rates without considering chromosomal change using MiSSE, we found that, for extant species, these rates do not appear to vary much (Fig. 3). Consequently, our reconstruction of chromosomal change across the deeper *Erebia* tree is unlikely to have been influenced by “hidden” speciation.

Here, we employed state-of-the-art phylogenomic models in one of the most karyologically diverse groups of butterflies, providing evidence for a macroevolutionary impact of major chromosomal rearrangements that equally occur in many other animal [*e.g*. ^48,60^] and plant [^49,50^] groups. Overall, we provide evidence that speciation rates are higher with increased chromosomal changes. Similar inferences of the impacts of chromosomal rearrangements are often carried out at higher taxonomic levels or across vast evolutionary timescales [*e.g*. ^61,62^; but see ^48^], potentially masking more fine-scaled patterns in younger clades. Our study bridges between these former investigations and microevolutionary studies that focus on one species or compare sibling species [*e.g*.^11,59,62^] by demonstrating within-genus differences of chromosomal fusion- and fission-related speciation. We highlight that chromosomal speciation may be more relevant in clades with more diversity in chromosome numbers. In this genomic era, ever more high-quality reference genomes can be generated and used to build phylogenomic frameworks, which will enable us to further unravel the complexities of chromosomal evolution and speciation.

## Materials and methods

### Data collection

Adult specimens of 83 *Erebia* species were collected between 2009 and 2021 (Table S1). Bodies were stored either in ethanol at −20 °C (n = 53) with wings separated, or pinned at room temperature (n = 30). For the latter, the wings were cut and stored separately prior to DNA extraction. DNA was extracted from thorax tissue using a Qiagen Blood & Tissue Kit (Qiagen AG, Hombrechtikon, Switzerland) following the standard manufacturer’s protocol. Paired-end sequencing libraries were constructed at the Department of Biosystems Science and Engineering (DBSSE) of ETH Zürich in Basel, followed by sequencing on an Illumina NovaSeq 6000. Samples were sequenced on two S1 flow cells.

Demultiplexed raw sequence reads were processed using fastp [^63^], trimming poly-G tails. Retained reads were mapped to the chromosome-resolved *Erebia ligea* reference genome [^64^], using bwa v0.7.17 [^65^]. Average mapping coverage was 38% but varied among species, reflecting phylogenetic distance to the reference genome (Fig. S9). One species (*E. atramentaria*) was omitted due to very low coverage (2.5%). SAMtools v.1.13 [^66^] was then used to remove unmapped, unpaired, or duplicated reads. A *pileup* file for each sample was generated with BCFtools v.1.12 [^67^] *mpileup*, followed by variant calling in BCFtools *call* [^68^]. Individual VCF files were then merged and subsequently filtered to remove i.) non-biallelic SNPs, ii.) insertions and deletions and adjacent SNPs within 5bps, iii.) SNPs with quality score <30, iv.) SNPs with more than 80% missing data, v.) SNPs with depths <4 or >25, vi.) SNPs with minor allele frequencies (MAF) <0.03, and vii.) SNPs falling within repetitive parts of the genome as identified by RepeatMasker 4.0.9 [^69^]. This resulted in a dataset containing 1.87 million SNPs from 82 *Erebia* samples.

Outgroup data was taken from chromosome-resolved assemblies of other Nymphalid butterflies that were publicly available at the time of analysis: *Pararge aegeria* [^70^], *Aphantopus hyperantus* [^71^], *Aglais urticae* [^72^], *Vanessa atalanta* [^73^] and *V. cardui* [^73^]. To obtain single copy orthologs (SCO) among all assemblies, each assembly including *E. ligea* was annotated with WebAugustus [^74,75^] in two steps: First, gene prediction was run using the standard Augustus species parameters for *Heliconius melpomene*. Then, pairwise SCOs between *E. ligea* and all other species were identified with Orthofinder [^76^]. These SCOs were subsequently used for gene prediction in a second run of WebAugustus.

A total of 4’505 SCOs among *E. ligea* and our selected outgroups were identified, of which 2’920 SCOs with ≥20 SNPs were retained for downstream analyses. Exons of each SCO were concatenated into a single coding region to create 2’920 separate VCF files extracted from the *Erebia* dataset with BEDtools [^77^]. VCF files were then transposed into FASTA format, the same exons were extracted for each outgroup species and aligned to the *Erebia* sequence files. Each resulting gene sequence file was aligned with MAFFT v.7.467 [^78^].

### Phylogenomic analyses

Maximum likelihood (ML) gene trees were estimated in IQTREE2 [Nguyen et al. 2015^79^], which first estimates the optimal substitution model for a gene alignment through ModelFinder [Kalyaanamoorthy et al. 2017^80^]. Gene trees were estimated with ultrafast bootstrap approximation for 1’000 iterations. A species consensus tree was then inferred using coalescent methods in ASTRAL-III [Zhang et al. 2018^81^]. Prior to this analysis, branches with <10% bootstrap support were collapsed to improve the accuracy of ASTRAL-III [Zhang et al. 2018^81^]. The resulting species tree had a normalized quartet score of 0.64, indicating 64 percent agreement between gene tree topologies, and high overall support (Fig. S1).

The robustness of the phylogeny was further validated following [^25^], *i.e*. selecting loci based on phylogenetically informative parameters to reduce incongruency [^25,82,83^]. Five validation datasets were created, each containing the 600 gene trees that i.) had the highest average bootstrap in IQTREE, ii.) showed the highest clocklikeness, indicating how well a gene tree approaches an ultrametric tree formation, iii.) had the lowest CG *vs*. AT contents, iv.) had the highest proportion of parsimony informative sites, and v.) showed the lowest mutation saturation potential. Datasets i-iv were generated with AMAS [^84^]. The saturation potential was calculated through regression slopes, with lower mutational saturation potential indicating a lower degree of amino acid substitutions, and a lower sensitivity to differences in evolutionary rates [^83,85^]. Consensus topologies for each dataset based on IQTREE gene trees were generated with ASTRAL-III. The average normalized quartet score was 0.68 (range 0.61-0.77). Comparisons between topologies of the overall dataset and the validation topologies indicated only minor differences concerning relationships within clades. All clades used for further analyses were supported in all validation datasets, with exception of the highest clocklikeness dataset, where the *pluto* clade included two formerly unassigned species (Figs. S2-6). The phylogeny based on the overall dataset, hereafter referred to as the *All gene* topology, was therefore used in downstream analyses to constrain dating analyses using fossil calibration.

### Divergence time estimation

Most state-dependent birth-death (SSE) models are based on ultrametric trees, with time expressed either in relative branch length equivalents or with branch lengths representing absolute time. The latter allows for the expression of state-dependent rates of diversification in terms of Events/Species/Myr, and is therefore more straightforward to interpret. We inferred an ultrametric time tree based on the 56 genes with <1% missing data, for a total of 82’923 base pairs, with the topology constrained according to the *All gene* topology. We chose to minimize missing data for the dating analysis as it occurred primarily in outgroups, which could bias branch lengths and relationships among outgroups, which would in turn impact the calibrations, which were placed among the outgroups.

No fossils are known for *Erebia* and few for Satyrini overall. *Lethe corbieri* [^86^], which has previously been used for dating Satyrini phylogenies [^87^], was therefore selected. As the two outgroup species within Satyrini, *P. aegeria* and *A. hyperantus*, span the breadth of the clade, their split coincides with the age of Satyrini. Thus, the age of *L. corbieri* (25.0 Mya) was taken as a conservative minimal bound for the age of Satyrini, with the median age set at 41.8 Mya, following the inference of [^87^]. The root of the tree represents the age of Nymphalidae, which was previously estimated to be 69.4 Mya (59.0 – 80.2 Mya) [^88^], which was taken as the median age for this node. The minimum age of Nymphalidae was bounded using the fossil *Vanessa amerindica*, as its placement within *Vanessa* is debated [^89^], and we considered placing this fossil closer to the base of Nymphalidae to be the more conservative approach.

Calculations of the divergence times were carried out by node dating in MrBayes v. 3.2.7 [^90^]. PartitionFinder [^91^] was used to partition the dataset and select the per-partition best substitution models. The *invgamma* model for among-site rate variation was the best model for all partitions and we further allowed for integration over the full GTR model space. Hard constraints were placed on all nodes based on the *All gene* topology, with the clade Satyrini constrained to *Erebia*, *P. aegeria*, and *A. hyperantus* to estimate its node age based on an offset exponential prior with 25.0 Mya as minimum and 41.8 Mya as median, while 33.7 Mya and 69.4 Mya served as the minimum and median bounds of the root. A rooted birth-death process was used to inform the prior probability distribution of branch lengths, strategy of extant taxa was set to random, and sampling probability was set to 0.0145 (87 species out of approximately 6’000 Nymphalidae). MrBayes was run for 10’000’000 MCMC generations in two independent runs, with four chains each with temperature set to 0.3, and a burn-in of 50%. All model parameters converged to an effective sampling size (ESS) >200 and potential scale reduction factors approached 1.00.

The root and outgroups were subsequently pruned in R version 3.6.3 [^92^] using the *extractClade* function of *addTaxa* [^93^]. In order to allow for comparisons of chromosome-related diversification rates, the six clades of *Erebia* for which chromosome data was available for at least two species (*ligea*, *pronoe*, *medusa*, *pluto*, *epiphron*, and *tyndarus*), were similarly extracted from the overall time-calibrated tree with their internal branch lengths and node ages preserved.

### Inference of ancestral chromosome numbers

ChromoSSE [^27^] was used to jointly infer ancestral chromosome evolution and the phylogenetic birth-death process within the Bayesian framework of RevBayes [^94^]. Chromosome numbers are available for 47 out of 82 included *Erebia* species (Table S1). Species with unknown chromosome numbers were treated as missing data. The maximum chromosome number allowed for inference was set at 56, five more than the known maximum of n = 51 for *E. iranica* to allow for computationally optimal parameter exploration. As polyploidy has never been recorded in Lepidoptera [^95^], the polyploidisation and demipolyploidisation probabilities were set to zero for both the anagenetic and cladogenetic parts of the model. The frequency of each possible chromosome number at the root of *Erebia* was estimated alongside all other parameters, to ensure a more accurate assessment of the root number. For the overall analysis, taxon sampling probability (rho_bd) was set to 0.82 as our ultrametric tree included 82 species out of the approximately 100 recorded for *Erebia* [^21,96^]. Stochastic character mapping of ancestral chromosome numbers was added to the model, with sampling every 10 generations. The inference of ancestral states and the model parameters were run in two independent runs of 10’000 MCMC generations each, sampled every 10 generations, with a burn-in of 20%. Convergence of the runs was assessed visually in Tracer v.1.7.1 [^97^], and the posterior parameter spaces of the runs were combined prior to further analyses. Priors for anagenetic parameters were established from an exponential distribution centred on 10.0, while priors for cladogenetic parameters were calculated from the number of taxa involved and the root age of the phylogeny following the standard ChromoSSE analysis [^27^]. For our analyses of clades, the same model setup was used as above, with the maximum chromosome number adjusted to the maximum count within that clade +5, but analyses were carried out as single runs, with more generations (25’000 MCMC generations for the *tyndarus* clade, and between 50’000-100’000 MCMC generations for all other clades, always with a burnin of 50%) depending on the size of the clade and the maximum chromosome count, as these impact computational intensity. Rho_bd was set to the percentage of known extant *Erebia* species that was sampled for each clade (between 0.75 and 1.00), the details of each parameter and run are recorded in Table S5. The convergence of each chain in each clade was assessed in Tracer v.1.7.1 [^97^], ensuring an ESS >200 for all parameters.

In order to validate the results of ChromoSSE and assess the impact of modelling the birth-death process along with chromosome evolution, a similar analysis was carried out in ChromEVOL [^31^] as modified according to [^27^], allowing to model cladogenetic changes without extinction. Parameter and burnin setup were similar to the ChromoSSE setup (Table S5).

### MiSSE

As SSE models have been suggested to suffer from excess false positives [^28^], we estimated speciation and extinction rates across the *Erebia* tree independently of chromosome states using MiSSE [^29^], which is based on the “hidden-states” state-dependent model HiSSE [^30^] and implemented in R. HiSSE seeks to overcome the common critique of SSE models of increased false-positive associations between diversification rates and a studied trait, by using hidden-Markov models to allow for hidden states, which are not sampled at the tips, alongside the studied focal traits. MiSSE instead considers only the hidden states themselves [^29^], allowing for a more accurate estimation of “background” shifts in diversification and extinction rates across given phylogeny compared to previous methods [^29^]. MiSSE was run with the greedy algorithm to determine an optimal set of hidden state models based on AIC, allowing four hidden states and 28 possible model combinations, with the sampling fraction set to 0.82 as in ChromoSSE. MiSSE was directed to stop model fitting when ΔAIC became < 2 (Table S6). Tip rates were then estimated after model averaging (summarized in Table S2) and the current speciation and extinction rates for extant species were compared among clades using ANOVAs.

### Model assessment and clade comparisons

In order to compare the estimated chromosomal parameters between clades, posterior parameter means were first computed from the burned-in posterior parameter spaces of the ChromoSSE and ChromEvol analyses separately, along with the 95% highest posterior densities (95%HPD) of each parameter. Posterior parameter distributions from ChromoSSE were then compared among clades using an ANOVA, followed by Tukey *post hoc* tests.

We assessed the difference in model fit between ChromoSSE and ChromEvol models for the overall *Erebia* tree and for each clade by calculating Bayes Factors and interpreting their size following [^98^]. As these models calculate most of the same parameters for the same phylogeny, we further quantified the differences between posterior distribution spaces of these models using *t*-tests. However, as ChromEvol estimates cladogenetic parameters as proportions of the total amount of cladogenetic change, cladogenetic parameters from the ChromoSSE analysis were first divided by the mean of the total speciation rate, thereby obtaining standardized parameters, allowing us to compare relative Events/Species/Myr.

## Acknowledgements

We thank Cristophe Praz for his advice on validation of phylogenetic inferences and Anatoly Filippov, Peter Jakubek for their aid in specimen collection. HA was supported by the Swiss National Science Foundation grant 310030_184934 *Genomic rearrangements and the origin of species* awarded to KL. JMdV was supported in part by Swiss National Science Foundation grant 310030_185251. RV was supported by project PID2019-107078GB-I00/ MCIN/AEI/10.13039/501100011033. VD was supported by the Academy of Finland (Academy Research Fellow, decisions no. 324988 and 352652). KO was supported by JSPS KAKENHI Grant Number JP21K15165. Analyses were performed at sciCORE (http://scicore.unibas.ch/) the scientific computing centre at University of Basel. The authors declare no conflict of interest.

## Author contributions

HA and KL conceived the project. RV, VD, KO, MC, TC and GI collected samples. HA carried out the analyses, supervised by KL and assisted by SK, JdV and LB. HA wrote the manuscript with input from KL, and all authors read and commented on the manuscript.

